# Space-number association in zebrafish

**DOI:** 10.64898/2026.03.04.709503

**Authors:** Davide Potrich, Mirko Zanon, Rosa Rugani, Valeria Anna Sovrano, Giorgio Vallortigara

## Abstract

Spatial-numerical associations reflect the tendency to map small numerosities to the left and larger numerosities to the right. Although widely documented in humans and some non-human species, its presence in teleost fish has remained unclear. Here, we investigated whether zebrafish exhibit a spatial-numerical association by presenting fish with a left–right numerical discrimination task in a controlled behavioral assay. Zebrafish showed a reliable leftward preference when selecting smaller numerosities and a rightward preference when selecting larger ones, indicating a systematic coupling between numerical magnitude and spatial direction. These results provide the first evidence of a number-space association in zebrafish and demonstrate that number–space mapping extends to a basal vertebrate lineage. This establishes zebrafish as a tractable model for probing the neurobiological foundations of number-space associations.

**Highlights:**

- Zebrafish map numerical magnitudes onto left–right spatial positions.
- Spatial biases are more robust for small numerosities but still present for larger ones.
- Continuous physical controls disproportionately affect the strength of spatial mapping for large sets.

## Introduction

Spatial-numerical associations (SNA) describe a consistent behavioral tendency to map numerical magnitude onto space. In humans, smaller numbers bias responses leftward and larger numbers rightward, a pattern commonly interpreted as evidence for a spatially organized mental number line. Although first identified in adults (starting from Galton’s original observations ^1^ and later formalized by Dehaene ^2^ as the SNARC effect), similar directional biases have since been documented across early development ^3^, and in populations with minimal formal education ^4^, indicating that space-number mapping does not depend on symbolic mathematics or cultural conventions (for a review, see ^5^).

Importantly, numerical spatialization is not uniquely human. Converging evidence from primates, birds, and even invertebrates shows that non-human species associate numerical magnitude with space ^6–8^. Leftward biases for smaller numerosities and rightward biases for larger ones have been reported in domestic chicks and Clark’s nutcrackers ^9,10^, rhesus macaques ^11,12^, and chimpanzees ^13^; even honeybees display comparable space-magnitude mapping, despite their radically different neural architecture^14,15^.

However, the universality of this mapping across all numerical ranges remains debated. Although the overall pattern of small-left and large-right associations appears broadly conserved in birds and mammals, individual variability in directional preference has been documented in primates and birds ^16–18^, and contrasting evidence exists in other contexts ^19,20^.

Two studies, respectively with domestic chicks and cleaner fish, provided a paradigmatic example ^10,20^. The seminal study on relative SNA in animals was conducted on domestic chicks, an animal model known to master a wide range of numerical abilities. During training chicks learnt to circumvent a panel displaying a target numerosity (5). At test they faced two panels, one on the left and one on the right, depicting the same numerosity, either smaller (2) or larger (8) than the target. The animals showed a robust leftward bias when presented with the smaller numerosity and a rightward bias when presented with the larger one ^10^. This indicates that the numerosity experienced during training functions as a reference magnitude stored in memory, against which future numerosities are compared. This aligns with evidence that zebrafish not only learn and remember visual information ^21,22^ but also exhibit representational abilities, such as maintaining a goal-object in memory after limited exposure ^23,24^. Yet, when a similar experimental paradigm was applied to cleaner fish (*Labroides dimidiatus*), no systematic numerical spatialization was observed ^20^. This failure might depend on the numerosities used, even if fish succeeded in a numerical discrimination task requiring them to distinguish between two simultaneously visible quantities. Another possible explanation can be temporal stability of numerical information in working memory, as supported by evidence that cleaner wrasse fails in both visual and spatial working memory tasks ^25^, indicating a bottleneck in maintaining representations over time. By contrast, the paradigm was originally designed for chicks, taking into account their working□memory duration and the specific experimental constraints ^10^, and using magnitudes that chicks had already been shown to discriminate successfully ^26,27^.

Crucially, numerical cognition is thought to rely on two distinct non-symbolic systems: the *Approximate Number System* (ANS), which represents large magnitudes noisily and is ratio-dependent, and the *Object Tracking System* (OTS), which allows for the precise tracking of small sets (typically ≤4 items, also referred to as *subitizing* range) ^28,29^. Notably, the capacity limit separating ‘small’ from ‘large’ numerosities is not absolute, but varies as a function of species, age, and task complexity ^28,30,31^ . In teleost fish, the picture seems particularly complex, often revealing dissociations between these systems. For instance, mosquitofish (*Gambusia holbrooki*) show precise discrimination for small sets (1 vs. 2 up to 3 vs. 4) but fail at the 4 vs. 5 transition, requiring a larger ratio (1:2, e.g., 4 vs. 8) to discriminate larger magnitudes ^32^. Similarly, guppies (*Poecilia reticulata*) successfully compare quantities within the small (3 vs. 4) or large (5 vs. 10) ranges but fail specifically when comparisons closely span the system boundary (3 vs. 5), recovering performance only when the numerical distance is increased (e.g., 3 vs. 6)^33^. In contrast, other species such as angelfish (*Pterophyllum scalare*) appear to follow a more continuous ratio-dependent limit (Weber’s law) without a sharp discontinuity for small numbers ^34^. In zebrafish (*Danio rerio*), quantity discrimination is well-established across both small and large ranges ^35^, supported by specific neurons in the dorso-central pallium that respond to numerical change ^36,37^. Indeed, by comparing these previous findings in parallel with the diversity of experimental paradigms used, it acts as a compelling suggestion that the specific involvement of the OTS versus the ANS is not rigid, but may crucially depend on task-specific demands and the cognitive load imposed by the experimental setting. This heterogeneity suggests that spatial-numerical mapping in fish may not be a generalized trait, but rather one that depends on the specific cognitive system (exact vs. approximate) recruited.

The two studies mentioned above, respectively with domestic chicks and cleaner fish, provided a paradigmatic example ^10,20^. Chicks were trained to circumvent a panel displaying a target numerosity (5) and subsequently tested with panels depicting either a smaller (2) or larger (8) numerosity. This design explicitly crosses the boundary between the small number range (subitizing/OTS) and the large number range (ANS). The animals showed a robust leftward bias when presented with the smaller numerosity and a rightward bias when presented with the larger one^10^. Yet, when a similar experimental paradigm was applied to cleaner fish (*Labroides dimidiatus*), which succeeded in a numerical discrimination task, no systematic numerical spatialization was observed ^20^. This suggests that while numerical discrimination is present, the spatial mapping of these quantities might depend on how the species processes distinct numerical ranges (small vs. large) or the temporal stability of their working memory for numerical versus spatial information. This is further supported by evidence that cleaner wrasse fish struggle with the early stages of both visual and spatial working memory paradigms ^25^, suggesting a bottleneck in maintaining representations over time.

Two main evolutionary scenarios remain plausible. The presence of SNA in taxa that diverged more than 500 million years ago —such as honey bees and vertebrates—suggests that this mapping between number and space may represent an ancient, conserved cognitive mechanism. Alternatively, SNA may have arisen independently in different lineages through convergent evolution, as an adaptive response to shared ecological demands. The heterogeneous findings in fish further complicate this picture: if the absence of SNA in some aquatic species is confirmed, it may indicate either a later emergence of the trait within specific vertebrate clades or a secondary loss in certain environments. Determining whether SNA reflects a deep ancestral heritage or a convergent cognitive adaptation remains an open challenge for comparative research ^38^. At the cognitive level, these findings suggest that SNA may arise from evolutionarily ancient mechanisms, but its expression may differ based on the cognitive architecture used to process numerosities, such as the capacity and duration of working memory required to bridge the stimulus-to-response interval. The robustness of the chick paradigm, which has been successfully applied in other species ^3,14,39^, combined with the informative null result in cleaner fish underscores the need for systematic investigations in model organisms like zebrafish to disentangle these factors. Zebrafish have already demonstrated the capacity to use numerical information in ordinal counting tasks ^40^ and proto-arithmetic contexts ^41^, while showing efficient one-trial memory processing ^42^.

In this study, we investigated whether zebrafish exhibit a SNA bias and whether this bias holds consistent when crossing the cognitive boundary between small and large numbers. We adapted a controlled behavioral task in which fish were required to discriminate between numerosity cues presented laterally (see Figure 1). Specifically, fish were initially trained to discriminate between a stimulus depicting a fixed numerosity (Experiment 1: five squares; Experiment 2: two or eight squares) and a blank stimulus. The choice of these specific numerosities (2, 5, and 8) allowed us to test the subjects’ responses to magnitudes that fall within the distinct cognitive ranges typically associated with the OTS and the ANS, while keeping a framework directly comparable with previous works ^10,20^.

**Figure 1.**
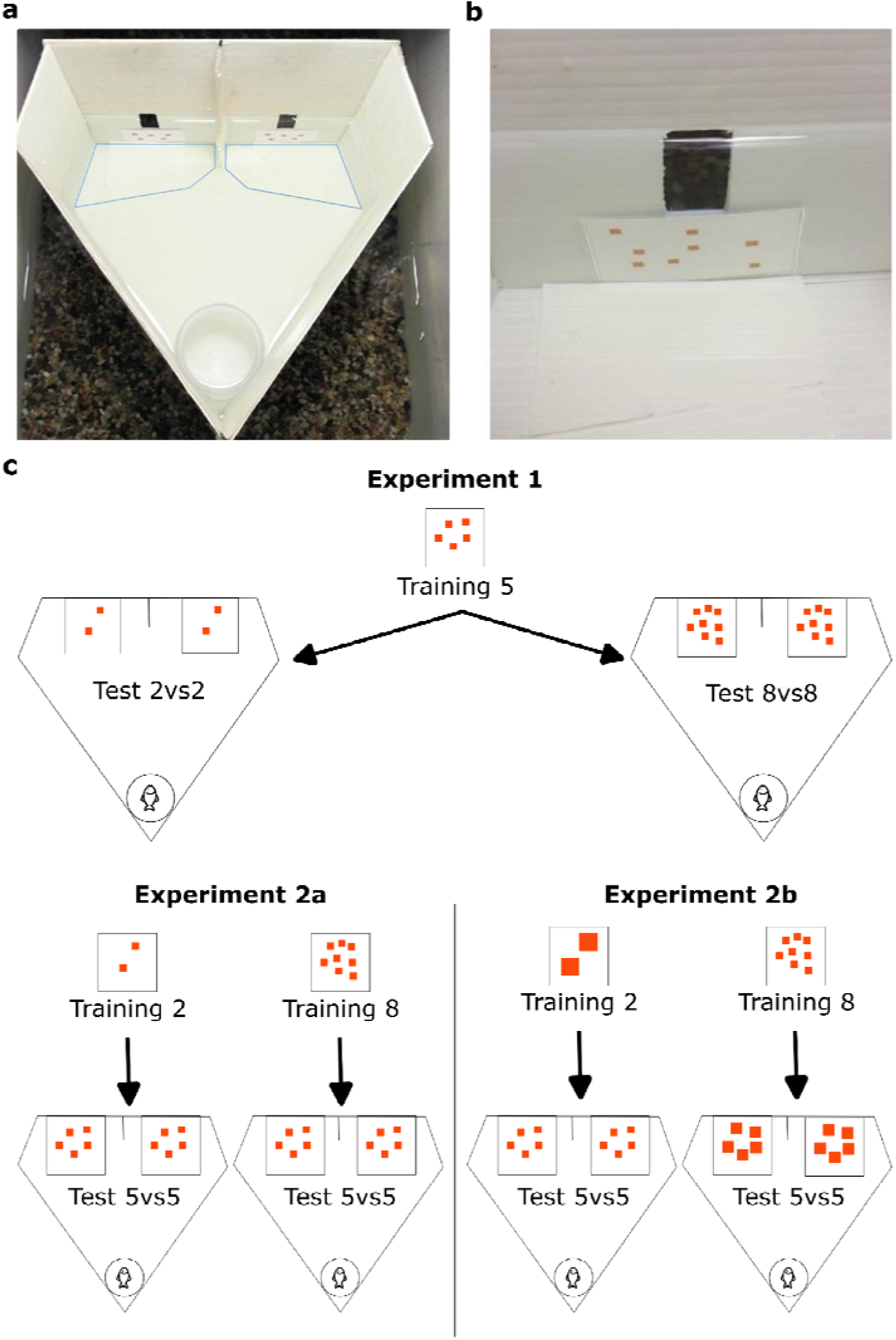
Apparatus and experiments schematic. **a)** Image of the apparatus. Decision areas are highlighted in blue. **b)** Image of the apparatus exit door with an example of the numerosity stimulus. **c)** Schematic of the two main experiments: Experiment 1 consists of a training with 5 squares followed by two tests (2vs2 squares and 8vs8), while Experiment 2 consisted of two groups, one trained with 2 squares and tested with 5vs5, and one trained with 8 squares and tested with 5vs5. In Experiments 1 and 2a, all squares have the same dimensions, while in Experiment 2b, the squares’ overall perimeter is preserved between training and test stimuli.

If fish associate numerosity with space across these ranges, in Experiment 1, they would select the left stimulus in the 2 vs. 2 test and the right one in the 8 vs. 8 test. If their spatial-numerical association is relative, in Experiment 2a, facing the same 5 vs. 5 test, we expect them to approach the left stimulus after training with 8 and the right one after training with 2. Finally, Experiment 2b controlled for physical variables (continuous quantities) to assess whether performance was driven by non-numerical perceptual cues.

A second hypothesis is that SNA in fish may be restricted to the small□number range or to large numerosities within large discernible numerical ratios. Under this account, our design should still be able to detect it: in Experiment 1, fish would be expected to choose the left stimulus in the 2 vs. 2 test after training with 5. The design also allows us to test the relativity of SNA: in Experiment 2, following training with 2, fish should preferentially select the right stimulus in the 5 vs. 5 comparison. Experiment 2b further evaluates whether such choices persist when continuous physical variables are controlled for. A delimitation to the small□number range would be supported by chance□level performance whenever 8 —falling within the large□number range— is used.

## Results

Fish successfully reached the learning criterion after an average of 99 ± 16 trials for Experiment 1, 172 ± 21 trials for Experiment 2, and 84 ± 10 trials for Experiment 2b. Test results for the different experiments are reported in Figure 2.

**Figure 2.**
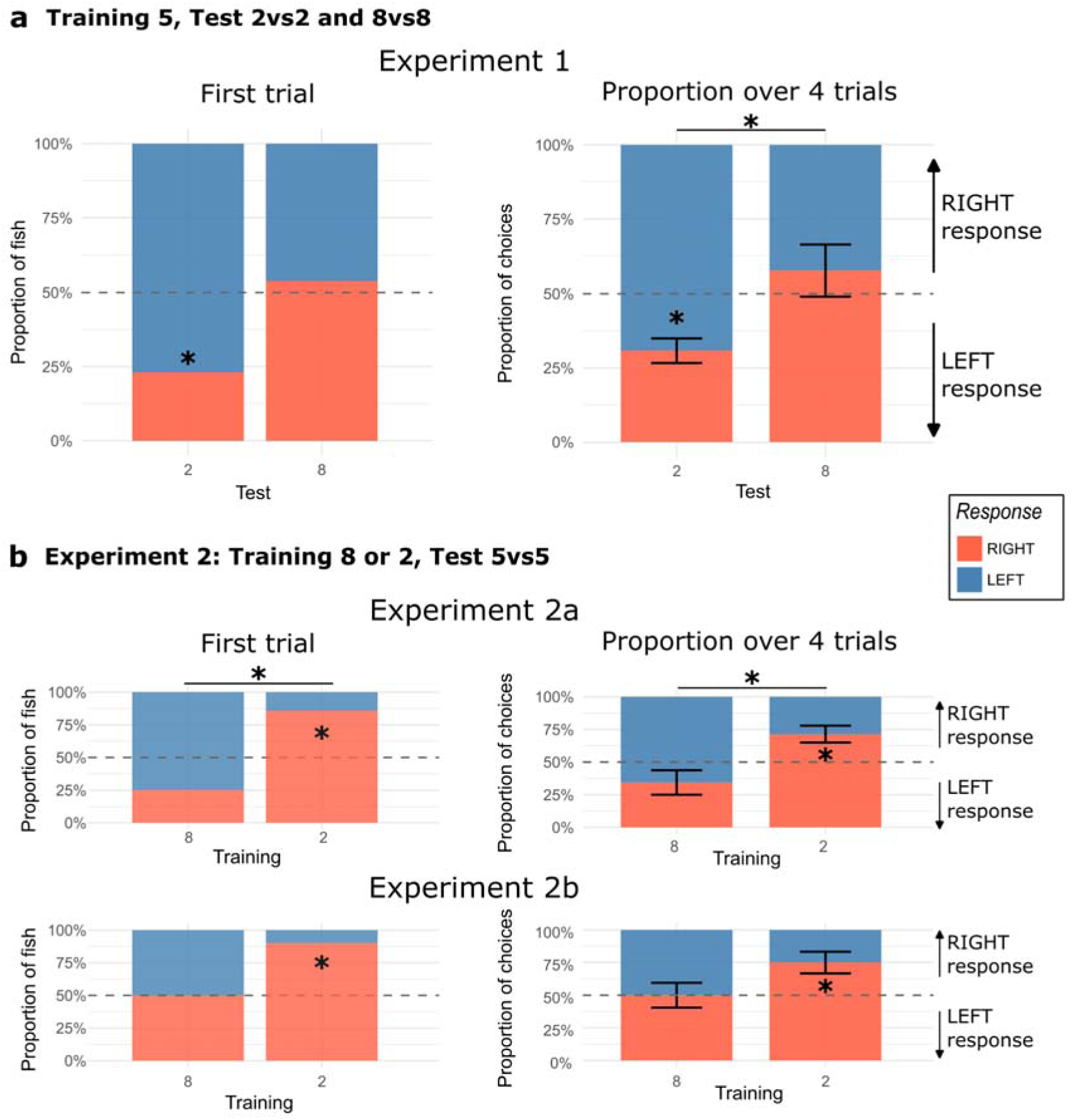
Experimental test results. Proportion of fish for first trial choice (left, lighter bars) and proportion of choices over the 4 test trials (right, darker bars), for the different experiments. **a)** Data for Experiment 1, with Training numerosity 5 and Test 2+8 (within subjects); **b)** Data for Experiment 2, with Training numerosity 2 or 8 (respectively, Experiment 2a and 2b; between subjects) and Test 5 (both for no perimeter control -top-and perimeter controlled stimuli -bottom-). * indicates p-values < 0.05 (binomial and Wilcoxon tests against 50% chance, paired Wilcoxon, Mann-Whitney U and exact Fisher tests for group comparisons).

### Asymmetric spatial biases for small versus large magnitudes

In Experiment 1, fish were trained on 5 squares and then presented with either a smaller (2 vs. 2) or larger (8 vs. 8) numerosity in separate test sessions. When presented with 2 vs. 2 squares, fish showed a significant leftward bias across test trials (proportion choosing left = 0.692 [0.611, 0.774], Cohen’s d = −1.284, p = 0.005, power = 0.989), although first-trial choices showed only a trend in the same direction (proportion = 0.769 [0.462, 0.950], Cohen’s h = −0.569, p = 0.093, power = 0.536). When presented with 8 vs. 8 squares, fish did not show a significant rightward bias across all trials (proportion choosing right = 0.577 [0.407, 0.747], Cohen’s d = 0.246, p = 0.389, power = 0.129) or on first trials (proportion = 0.538 [0.251, 0.808], Cohen’s h = 0.077, p = 1.000, power = 0.059), although the direction of the response was consistent with SNA predictions. Critically, direct comparison between the two test conditions revealed that fish exhibited significantly different lateral biases when presented with smaller versus larger numerosities across all trials (paired Wilcoxon test: difference = −0.269 [−0.480, −0.058], Cohen’s d (paired) = −0.694, p = 0.043, power = 0.632), with first trial performance showing a similar pattern (Fisher’s exact test: difference = −0.308 [−0.663, 0.047], Cohen’s g = 2.000, p = 0.192, power = 0.644).

### Stronger spatial mapping following training with small numerosities

In Experiment 2, separate groups of fish were trained on either 2 or 8 squares and then presented with 5 vs. 5 squares at test. Fish trained with 2 squares (when presented with a larger numerosity at test) showed a significant rightward bias across all test trials with 5 squares (proportion choosing right = 0.714 [0.586, 0.842], Cohen’s d = 1.242, p = 0.048, power = 0.781), with first trials performance showing a trend in the same direction (proportion = 0.857 [0.421, 0.996], Cohen’s h = 0.796, p = 0.125, power = 0.558). Fish trained with 8 squares (when presented with a smaller numerosity at test) did not show a significant leftward bias across all trials (proportion choosing left = 0.656 [0.473, 0.840], Cohen’s d = −0.589, p = 0.168, power = 0.303) or on first trials (proportion = 0.750 [0.349, 0.968], Cohen’s h = −0.524, p = 0.289, power = 0.316), though choices were once again directionally consistent with SNA expectations. Between-group comparison demonstrated that fish trained with different numerosity exhibited significantly different spatial responses when experiencing the same intermediate test numerosity across all trials (Mann-Whitney U: difference = 0.371 [0.122, 0.619], Cohen’s d = 1.631, p = 0.014, power = 0.830), with first trials also showing a significant difference (Fisher’s exact test: difference = 0.607 [0.211, 1.004], Cohen’s h = 1.319, p = 0.041, power = 0.722).

### Asymmetric robustness of small versus large magnitude mapping to continuous variable controls

In Experiment 2b (experiment controlling for continuous variables), fish trained on 2 squares maintained a significant SNA-consistent rightward bias when presented with 5 squares across all test trials (proportion choosing right = 0.750 [0.587, 0.913], Cohen’s d = 0.949, p = 0.031, power = 0.761) and on first test trials (proportion = 0.900 [0.555, 0.997], Cohen’s h = 0.927, p = 0.022, power = 0.835). In contrast, fish trained on 8 squares exhibited no lateral bias, with performance dropping to chance level across all trials (proportion choosing left = 0.500 [0.315, 0.685], Cohen’s d = 0.000, p = 1.000, power = 0.050) and on first trials (proportion = 0.500 [0.157, 0.843], Cohen’s h = 0.000, p = 1.000, power = 0.050), indicating no numerical spatialization. Between-group comparison revealed that fish trained with different numerosity showed a trend toward different spatial responses across all trials (Mann-Whitney U: difference = −0.250 [−0.518, 0.018], Cohen’s d = −0.943, p = 0.074, power = 0.463), with first trials showing a similar pattern (Fisher’s exact test: difference = −0.400 [−0.793, −0.007], Cohen’s h = −0.927, p = 0.118, power = 0.498).

## Discussion

The present study provides compelling evidence of spatial-numerical association (SNA) in zebrafish, demonstrating that this species spontaneously maps numerical magnitudes onto spatial left-right positions in a manner consistent with the *mental number line* observed in humans and other vertebrates ^6^. Fish trained on a reference numerosity and then presented with smaller or larger numerosity showed systematic lateral biases, with smaller numerosities leading to leftward choices and larger numerosity to rightward choices. This finding extends the phylogenetic distribution of the spatial numerical association beyond mammals and birds, suggesting that SNA represents a fundamental organizing principle of numerical cognition shared across vertebrates. Moreover, this extends the study of space-number mapping to a genetically tractable vertebrate model, providing a powerful framework for investigating the neural and evolutionary foundations of numerical cognition.

Our results reveal intriguing asymmetries in the strength of SNA depending on training numerosity, which may reflect distinct processing mechanisms for small versus large magnitudes. Fish trained with small numerosity (2 or 5) consistently exhibited robust spatial biases across test conditions. However, associations involving the larger numerosity (8) were less reliable: training with 5 and testing with 8 produced directionally consistent but non-significant biases, while training with 8 and testing with 5 yielded only marginally significant effects. This pattern aligns with recent evidence highlighting the role of absolute magnitude in SNA. As noted by Roth et al. (2025) ^43^, lower numerical ranges tend to produce stronger SNA, where both relative and absolute small numbers are associated with leftward space, while only relatively large numbers are linked to rightward space. Consequently, the weaker effects observed with larger sets may result from the decreased precision of the Approximate Number System (ANS; consistent with Weber’s law), whereas the robust performance with small sets may be supported by the precise Object Tracking System (OTS). Indeed, the 5 vs. 8 contrast (ratio 0.625) sits immediately adjacent to the known limit of zebrafish numerical precision. Previous work demonstrates that while zebrafish can discriminate at a 0.67 ratio (4 vs. 6), they fail at 0.75 (6 vs. 8) and even struggle with the 3 vs. 4 small-number limit ^35^. This suggests that the 5 vs. 8 comparison is inherently demanding; the resulting noisy internal representation likely destabilizes the spatial-numerical mapping compared to the high-precision OTS used for smaller sets. This destabilization may be further exacerbated by limits in visual memory. While zebrafish exhibit efficient one-trial memory in simpler contexts ^42^, the cognitive load of maintaining a representation of ‘8’ might lead to a comparison drawback during the decision-making process.

The distinction between these cognitive systems is further supported by the findings in the perimeter-balanced condition. Fish trained on 2 squares maintained robust SNA biases when presented with 5 squares at test, even when perimeter cues were controlled. In stark contrast, fish trained on 8 squares showed no lateral bias, with performance dropping to chance levels. This asymmetry suggests that fish trained with larger numerosity (relying on the ANS) may rely more strongly on continuous magnitude cues during learning, and when these cues are dissociated from numerosity at test, the resulting conflict disrupts spatial mapping. Conversely, the robust performance with small numbers suggests that the OTS may be more resilient to continuous physical variations, focusing instead on discrete item identity ^28,29,36^.

When compared with existing evidence, our results highlight a contrast between the present study and the findings reported by Triki and Bshary ^20^ in the cleaner fish *Labroides dimidiatus*. Specifically, *L. dimidiatus* showed no evidence of a spatial–numerical association, despite using similar numerosities. This discrepancy might be attributable to ecological factors or methodological differences that impact attentional engagement. *L. dimidiatus* is a marine species, whereas zebrafish are freshwater fish, which may favor different spatial processing strategies and capacities for spatial working and reference memory ^22^. A key difference also pertains to the experimental paradigm. In Triki and Bshary’s procedure^20^, cleaner fish had to swim around a central stimulus, a design that may have allowed subjects to solve the task without actively attending to numerosity. In our paradigm, by contrast, attentional engagement with the numerical stimulus was deliberately emphasized, as reinforcement was contingent solely on selecting the panel with the elements. This aspect is especially relevant for zebrafish, which require explicit task relevance to direct attention to specific stimulus features ^44^. Future comparative work should clarify whether the system for small numbers (as seen in *Gambusia*) is universally linked to spatial mapping, or if species-specific ecological constraints decouple these systems. A more fundamental explanation for this discrepancy may lie in inherent differences in working memory capacity. Recent comparative studies indicate that cleaner fish (*L. dimidiatus*) perform poorly in both visual and spatial working memory paradigms ^25^, potentially explaining their failure to exhibit SNA. In contrast, zebrafish possess a more robust memory architecture, enabling them to integrate numerical and spatial information ^40^. This suggests that a minimum threshold of working memory stability is a prerequisite for the emergence of SNA.

In conclusion, the present findings establish zebrafish as a tractable model for investigating the biological mechanisms of spatial-numerical association and pave the way for future research. The demonstration of SNA in zebrafish is particularly significant given the species’ status as a leading genetic model organism ^45^. The extensive genetic toolkit available for zebrafish, including CRISPR/Cas9 gene editing, transgenic lines with fluorescently labeled neurons, and well-characterized neural circuits, positions this species as an ideal system for dissecting the biological foundations of numerical cognition ^37,46–48^. Developmental studies could determine whether these mappings are innate or shaped by specific experiences. Genetic approaches could help identify candidate genes and neural pathways via targeted knockouts or mutant screening. Calcium imaging could localize brain regions mediating these effects, potentially revealing whether homologous circuits underlie the mental number line across vertebrates. Future work could also test broader ranges of numerosity and systematically manipulate continuous variables to determine their contributions to spatial choices. Additionally, studies could examine individual differences in spatial biases and investigate whether training can modulate spontaneous spatial-numerical associations.

By bridging comparative psychology and molecular neuroscience, zebrafish research can provide unique insights into how abstract numerical information is represented and spatially organized in the mind. The growing body of evidence for SNA across diverse species (including humans, non-human primates, birds, and now fish) suggests that spatial-numerical associations constitute a deeply conserved feature of vertebrate cognition. Zebrafish, with their unique combination of demonstrable numerical abilities and powerful genetic and neurobiological tools, provide an unprecedented opportunity to investigate the mechanisms underlying this phenomenon, potentially shedding light on the evolution and development of numerical cognition across species.

### Limitations of the study

While our results provide the first evidence of bilateral SNA in zebrafish, several limitations must be considered. First, the sample size in the experiments was relatively small, which may have reduced statistical power. Nevertheless, we obtained strong a posteriori effect sizes when small numerosities were used, showing a reliable association of a numerosity smaller than the training one with the left space, and of a larger one with the right space, when the numerical range fell within the limits of the Object Tracking System (2–5). For the larger numerosity comparisons, moderate/small effect sizes are possibly explained by a natural variance that is higher due to Weber’s law. Future studies can also better highlight possible individual differences ^16–18^ (see individual scores in Supplementary Figure 1).

Regarding continuous physical variables, our control condition (Experiment 2b) was limited to equating the total perimeter of the stimuli. This manipulation effectively made the total surface area inversely proportional to numerosity, thus putting area cues in contrast with numerical cues. However, we did not systematically control for other non-numerical variables such as density or convex hull. While the robustness of the “small number” results suggests that zebrafish can process discrete item identity despite these variations, the disruption of performance in the “large number” condition suggests that these continuous magnitudes may play a stronger role when the ANS is recruited. Future studies should employ multi-variable controls to disentangle the specific contributions of density, area, and convex hull across a wider range of numerosities.

Finally, the current study tested a limited set of numerosities (2, 5, and 8). To fully map the boundary between the OTS and the ANS in zebrafish, future work should employ a broader range of values (e.g., involving numerosities 3,4 and also bigger than 8) to precisely identify the point at which the spatial mapping degrades or shifts strategy. Moreover, the transition from training to test imposes a specific cognitive load. In Experiment 2 (the 5 vs 5 test), the fish must retrieve a reference value from memory and compare it to the current stimulus. If this comparison exceeds the species’ processing limit (as suggested by the lack of success in other teleosts with poor working memory ^25^) the spatial-numerical mapping may fail.

## Methods

### EXPERIMENTAL MODEL AND STUDY PARTICIPANT DETAILS

In the experiments, adult male zebrafish (*Danio rerio*) served as test subjects, while female zebrafish were used as a social reward. Fish were provided by a local pet shop (“Acquario G” di Segatta Stefano, Trento, Italy) housed in 3.5 L plastic tanks within an automated aquarium system (ZebTEC Benchtop, Tecniplast), in same-sex groups of 10 individuals. Water temperature was maintained at 26 °C, with a 12:12 - h LD cycle. Animals were fed dry food (Sera GVG-Mix) three times per day.

During the experimental period, subjects were individually housed in 20-L plastic aquaria (23 × 38 × 25 cm) containing gravel and artificial vegetation to simulate a naturalistic environment. Although physically isolated, fish had visual access to conspecifics through a mesh divider, minimizing potential effects of social deprivation. Total body length ranged from 4 to 5 cm across subjects.

Fifteen fish were observed in Experiment 1, of which 13 reached the learning criterion and subsequently proceeded to the test phase. Experiment 2a involved 20 fish, of which 15 reached the criterion (7 trained with 2 elements and 8 trained with 8 elements). Experiment 2b involved 20 fish, of which 18 fish reached the criterion (10 trained with 2 elements and 8 trained with 8 elements).

All experimental procedures complied with current Italian and European Union legislation regarding the ethical use of animals in research. The study was approved and licensed by the Italian Ministry of Health (auth. no. 1111/15-PR, prot. no. 13/2015 and auth. no. 893/2018-PR).

### METHOD DETAILS

#### Apparatus and stimuli

The experimental setup consisted of a diamond-shaped arena (28 × 25 × 19 cm) made of white plastic (Poliplak; see Figure 1a). Two identical rectangular exit openings (1.5 × 4.5 cm) were symmetrically positioned on the longest wall of the arena, 4.5 cm above the bottom. These openings allowed access from the central arena to an external compartment.

The arena was placed inside a larger opaque tank (35 × 49 × 27 cm) containing gravel, artificial plants, and two female conspecifics, which provided a strong social incentive for subjects to exit the arena. Directly below each exit opening, a removable white plastic panel displayed a fixed number of orange squares serving as the numerical stimuli (see Figure 1b). A vertical plastic divider protruded between the two exits, separating the arena into left and right choice zones.

Water temperature was kept at 26 °C and water was circulated using a pump and filtration system (Micro Jet Filter MCF 40). Illumination was provided by a 60 W fluorescent lamp, and fish behavior was recorded from above using a webcam (Microsoft LifeCam Studio) placed approximately 50 cm above the apparatus.

The stimuli consisted of two-dimensional orange (RGB: 252,72,11) squares printed on laminated white cards. For each stimulus, the size of the individual squares was held constant, while their spatial arrangement varied randomly (with a minimum inter-distance of 4 mm and maximum of 21 mm). For each numerosity, eight distinct spatial configurations were created. In Experiments 1 and 2, all squares had a side length of 3 mm . These squared elements, viewed from a distance of 17 cm (starting position-stimuli distance), subtend a visual angle of approximately 1°. Considering a minimum inter-stimulus gap of 3 mm, the resulting spatial frequency is approximately 0.5 cycles per degree (cpd). Because the actual inter-distance values were typically larger than 3 mm, which would result in an even lower and more easily resolvable spatial frequency, this represents a conservative estimate. This value remains below the reported visual acuity threshold for adult zebrafish (approximately 0.6 cpd ^49^), ensuring the stimuli were clearly resolvable already from the starting position. In Experiment 2b (experiment controlling for continuous variables), stimulus size ranged from 3 mm to 7.5 mm. In this latter condition, the continuous non-numerical variable controlled across different numerosity was the overall perimeter. Balancing the perimeter, allowed us to evaluate its role in the spatial association and to indirectly assess the influence of overall area. Specifically, when the perimeter was equalized between two arrays displaying different numerosity, the stimulus with fewer elements had a larger total area than the one with more elements ^50^. If the animal had relied on the overall area of the stimulus rather than the number, a reversed spatial numerical preference would have been observed.

#### Behavioural assay

The experiment consisted of a training phase followed by a test phase. During training, only one of the two exit doors was accessible, while the other was blocked using a transparent plastic sheet. The open door was indicated by a plastic card displaying a fixed number of elements, whereas a blank white card was positioned beneath the blocked door. Each fish underwent one daily session consisting of 10 trials. Before the start of each session, the subject was transferred from its home aquarium to the outer, enriched environment for a 5- min acclimation period. At the beginning of each trial, the fish was confined within a transparent plastic cylinder (6 cm diameter) positioned in front of the stimuli. After a 15-sec delay, the cylinder was raised, allowing the fish to explore the arena freely. A correction procedure was applied: if an incorrect door was approached, the fish was allowed to revise its choice until it successfully exited through the open door or until the maximum trial duration of 15-min had elapsed. A choice was recorded when the fish contacted a door with its snout (in the case of the blocked door) or passed through the open door. Reinforcement was contingent on performance, such that a correct first choice resulted in a 6-min access to food and social companions, whereas one or more incorrect choices resulted in a 3-min interval without food and conspecifics. If no choice was made within the allotted time, the fish was given a 5-min pause. Across trials, the side of the reinforced exit (left or right) and the spatial configuration of the elements were randomized to ensure that numerosity was the only reliable cue. Learning was considered achieved when a subject reached at least 70% correct choices in two consecutive sessions, after which it proceeded to the test phase.

In the test phase, subjects completed four unrewarded trials with both exit doors closed. The test phase consisted of four trials. In each trial, two identical stimuli depicting numerosity different from that used during training were placed beneath the left and right exits. Restricted choice areas were delineated in front of each door using white plastic panels on the tank floor, defining discrete left and right zones and minimizing a central neutral region (see Figure 1a; areas marked in blue). Before each trial, the fish was again confined in the cylinder and released after 15-sec; the first approach was recorded when the fish entered one of the zones with its entire body. Inter-trial intervals lasted 5-min, during which the fish could freely swim in the outer environment. For each subject, both the direction of the first approach and the total number of approaches across the four test trials were analyzed.

Two main experiments were conducted. Experiment 1 employed a within-subjects design in which fish were trained on a target numerosity of 5 squares and subsequently observed in tests with numerosity of 2 vs. 2 and 8 vs. 8 squares; the order of the two test conditions was counterbalanced across subjects. Experiment 2 used a between-subjects design in which fish were trained on either 2 or 8 elements and observed in the test with a numerosity of 5 vs. 5 squares. Experiment 2 was further divided into two variants: Experiment 2a, in which continuous quantitative properties covaried with numerosity, and Experiment 2b, in which stimuli were controlled for overall perimeter that remained constant across training and test stimuli.

### QUANTIFICATION AND STATISTICAL ANALYSIS

All analyses were performed separately for the two datasets using R software (version 2025.09.2+418; R Core Team, 2025).

In Experiment 1, each fish was observed in a within-subjects design across four trials per Test and two Test conditions. For each experiment, we summarized each fish’s behavior by calculating the proportion of right-side choices across the four trials within a given Test. These per-fish proportions were then used as independent observations at the group level. To determine whether fish exhibited a systematic side bias, the average proportion of right choices across all fish was compared against chance (50%) using a one-sample Wilcoxon signed-rank test (wilcox.test function with mu = 0.5, exact = FALSE). This non-parametric test was chosen because the data are proportions bounded between 0 and 1 and may not satisfy the normality assumptions required for a t-test. Moreover, because some fish had identical proportions across trials (ties), an approximate calculation of the p-value was used instead of the exact method, which cannot handle tied values. For first-trial data only, each fish contributes a single binary choice (left or right), which was tested against chance using an exact binomial test (binom.test function in R). To assess whether fish showed different side preferences between the two Test conditions, we performed paired-sample comparisons accounting for the within-subjects design. For aggregated trial data, we used a paired Wilcoxon signed-rank test (*wilcox.test* with paired = TRUE), which compares the distribution of paired differences without assuming normality. For first-trial data, we used McNemar’s test (mcnemar.test with correct = TRUE), which is specifically designed for paired categorical data and evaluates whether fish changed their choices between conditions more than expected by chance.

In Experiment 2, fish were assigned to different Training conditions in a between-subjects design, with each fish experiencing only one condition and being observed across four trials. Proportions of right-side choices were calculated for each fish within its assigned Training condition. Comparisons against chance (50%) were performed using the same one-sample Wilcoxon signed-rank test as in Experiment 1. For first-trial data, each fish’s binary choice was analyzed using an exact binomial test. In this between-subjects design, each fish’s proportion was treated as an independent observation, unlike the repeated-measures structure in Experiment 1. To compare side preferences between Training conditions, we used independent-samples tests. For aggregated trial data, we employed the Mann-Whitney U test (*wilcox.test* with paired = FALSE), a non-parametric test for comparing distributions between independent groups. For first-trial data, we used Fisher’s exact test (fisher.test) reported as a conservative alternative particularly appropriate for small sample sizes.

For both experiments, post-hoc statistical power was calculated using the *pwr* package with observed effect sizes and α = 0.05. For single-group comparisons, we used *pwr.t.test* as an approximation for Wilcoxon-tested comparisons and *pwr.p.test* for binomial tests (one-sample proportion tests). For paired comparisons (Experiment 1), we used *pwr.t.test*with type = “paired” for aggregated trial data and *pwr.p.test* for paired first-trial proportion data. For between-subjects comparisons (Experiment 2), we used *pwr.t2n.test* for aggregated trial data and *pwr.2p2n.test* for first-trial proportions. All tests were two-tailed.

Effect sizes were calculated using measures appropriate to each test type and design. For comparisons against chance in aggregated trial analyses (Wilcoxon tests), we computed an estimate of Cohen’s d as the standardized difference between the mean proportion and chance level: d = (M - 0.5) / SD, where M is the mean proportion and SD is the standard deviation across fish. For first-trial binomial tests, we calculated Cohen’s h = 2[arcsin(√p_obs) - arcsin(√0.5)], which is specifically designed for comparing proportions and uses an arcsine transformation to stabilize variance. For paired comparisons (Experiment 1), we calculated Cohen’s d for paired samples: d_paired = M_diff / SD_diff, where M_diff is the mean of the paired differences and SD_diff is the standard deviation of those differences. For McNemar’s test, we calculated Cohen’s g = (b - c) / √(b + c), where b and c represent the numbers of discordant pairs. For between-subjects comparisons (Experiment 2), we calculated Cohen’s d using the pooled standard deviation: d = (M_₁_ - M_₂_) / SD_pooled, where SD_pooled = √[((n_₁_-1)SD_₁_² + (n_₂_-1)SD_₂_²) / (n_₁_+n_₂_-2)], and Cohen’s h for independent proportions: h = 2[arcsin(√p_₁_) - arcsin(√p_₂_)]. All effect sizes were interpreted using conventional benchmarks: |d|, |h|, or |g| = 0.2 (small), 0.5 (medium), and 0.8 (large).

Confidence intervals (95%) were calculated to quantify estimation uncertainty. For single-group aggregated trial data, CIs were computed using the standard error of the mean proportion: CI = M ± 1.96 × (SD/√n). For first-trial analyses, exact Clopper-Pearson confidence intervals were obtained directly from the binomial test. For paired differences in aggregated trial data (Experiment 1), CIs were calculated as CI_diff = M_diff ± 1.96 × (SD_diff/√n). For between-subjects comparisons of aggregated trial data (Experiment 2), CIs for the mean difference were obtained using Welch’s t-test (t.test with var.equal = FALSE), which does not assume equal variances between groups. For first-trial comparisons in both datasets, CIs for the difference in proportions were calculated using a normal approximation with appropriate standard errors for paired or independent designs.

## Resource availability

### Lead contact

Further information and requests should be directed to and will be fulfilled by the corresponding author, Giorgio Vallortigara

### Materials availability

This study did not generate new unique reagents nor new animal lines.

### Data and code availability

- Data: Data is available in Figshare repository: https://doi.org/10.6084/m9.figshare.30869576
- Code: Code used for the analysis of this work is available in Figshare repository https://doi.org/10.6084/m9.figshare.30869576
- Additional information: Any additional information can be required to corresponding contact.

## Acknowledgments

This project was funded by the European Research Council (ERC) under the European Union’s Horizon 2020 research and innovation programme, Grant agreement No. 833504 (SPANUMBRA) to GV; by FARE–Ricerca in Italia: Framework per l’Attrazione e il Rafforzamento delle Eccellenze per la ricerca in Italia, III edition, project “NUMBRISH–The neurobiology of numerical cognition: searching for a molecular genetic signature in the zebrafish brain” Prot. R20YL9WN9N to GV; by PRIN - Progetti di Rilevante Interesse Nazionale, 2019 “Number-space association: A comparative developmental and neurobiological approach” to GV, VAS, and 2022 Grant Agreement No. P2022TKY7B (PNRR): “The emergence of proto-arithmetic abilities with empty and non-empty sets” to GV. This project has also received support from the Italian Ministry of University and Research through the Research Project of National Relevance (PRIN) - 2022 Prot. 202254RHRT, to RR. We wish thanking Nicola Corbellini, Chiara Franceschi, Francesca Morbioli, Anna Mutti, Alessia Panizza, Elisabetta Pedrocco, Valerio Rubino, Alessandra Venezia, and Johannes Waibl, for their partial assistance with the data collection; Ciro Petrone and Michela Maffei for the animal care and welfare.

## Authors’ contribution

Conceptualization: DP, GV, RR, VAS

Setup implementation: DP

Data analysis: MZ

Funding acquisition: GV, VAS

Supervision: GV, VAS, DP, RR

Writing original draft: DP, MZ.

All authors reviewed and edited the final version of the manuscript.

## Declaration of generative AI and AI-assisted technologies in the writing process

During the preparation of this work, the authors used ChatGPT exclusively to improve English and readability in some sentences. After using this tool, the authors reviewed and edited the content as needed and took full responsibility for the content of the publication.

